# Modulating Cardiac-Gut Microbiome Interaction Post-Myocardial Infarction with Engineered Bacteria

**DOI:** 10.1101/2025.07.10.664025

**Authors:** Tushar Madaan, Michelle L. Nieman, Tushar Rakheja, Nabil Siddiqui, Dominick Gherardini, Mathieu Sertorio, Nitin S. Kamble, Shindu C. Thomas, Alexandra Hartman, Sheryl E. Koch, Lance Knipper, Vito A. Marino, Sakthivel Sadayappan, John N. Lorenz, John P. Konhilas, Nalinikanth Kotagiri

## Abstract

The gut microbiome plays a critical role in the pathophysiology of acute myocardial infarction (MI). MI events significantly impact intestinal integrity which results in leakage of bacterial products into the systemic circulation. We demonstrate that MI not only compromises intestinal integrity, leading to systemic leakage of bacterial products like LPS, but also results in the translocation and colonization of live, intact gut bacteria in the MI heart – a novel aspect of the heart-gut axis. Our initial findings with natural murine gut microbiome were substantiated using orally administered E. coli Nissle 1917 (EcN), as a tracer bacterium. Furthermore, we engineered EcN to express the microbial anti-inflammatory molecule (MAM) derived from the probiotic *Faecalibacterium prausnitzii*. Treatment with this engineered strain, EcN-MAM, led to significantly improved survival and cardiac function in MI mice. This was attributed to enhanced gut barrier integrity, resulting in reduced systemic bacterial permeation and subsequent inflammation. These findings shed light on a previously unrecognized dimension of the heart-gut axis and highlight the potential of microbiome-based interventions in MI management.

## INTRODUCTION

Myocardial infarction (MI) represents a life-threatening cardiac emergency characterized by hypoxia, and subsequent myocardial necrosis^1–3^. Its pathophysiology involves complex mechanisms, including plaque rupture and thrombosis, often precipitated by unstable cardiac ischemia^4^. Globally, MI remains a leading cause of morbidity and mortality, exhibiting varying trends and prevalence across different demographics and regions^5,6^. While its global prevalence is 3.8% in individuals under 60, it rises to 9.5% in those over 60, highlighting its significant health burden^7^.

Recent advancements in cardiovascular research have underscored the gut-heart axis, particularly the role of the gut microbiome in the pathophysiology of MI^8^. The gut microbiome is increasingly recognized for its influence on overall cardiac health, with evidence suggesting that microbial imbalance may contribute to cardiovascular diseases^9,10^. Post-MI, compromised cardiac function and reduced intestinal perfusion lead to intestinal hyperpermeability, allowing gut microbiome leakage into the systemic circulation^11–13^. This leads to MI patients exhibiting elevated systemic microbial richness and diversity, which correlate with heightened levels of bacterial biomarkers such as lipopolysaccharides (LPS) and D-lactate^11^. These markers are associated with increased inflammation and subsequently reduced left ventricular function, crucially impacting patient survival and outcomes^11^.

Prior studies have demonstrated the ability of facultative anaerobic gut bacteria such as *E. coli* Nissle 1917 (EcN) to colonize and proliferate hypoxic regions of the body such as solid tumors post systemic introduction^14,15^. Obstruction of the normal cardiac blood flow due to MI creates a similar region of hypoxia and necrosis in the myocardium which can potentially be a homing ground for gut bacteria leaking into the systemic circulation after an MI attack^16,17^. Furthermore, persistent, long-term inflammation is a major cause for heart failure and progressive ventricular dilatation post-MI^18,19^. Systemic leakage of bacterial LPS can lead to activation of toll-like receptor 4 which can result in adverse post-infarct maladaptive left ventricular remodeling, cardiac fibrosis, and impaired cardiac function^20,21^. Clinical studies have demonstrated the significant influence of LPS on the cardiovascular system through inflammatory and immune pathways^11,22,23^.

Our study aims to bridge these research gaps by investigating the phenomenon of intestinal bacteria translocation and colonization in heart tissue post-MI. We also explore the clinical implications of our findings, potentially offering new therapeutic approaches for MI.

In light of the gut microbiome’s critical role in the pathophysiology of MI, attempts have been made to modulate the gut microbiota to help in the amelioration of MI^8^. Antibiotic-mediated gut microbiome abrogation has yielded mixed results. While one study reported that suppression of gut microbial translocation by depletion of gut bacteria using polymyxin B resulted in reduced infarct size, reduced local inflammation and subdued monocyte infiltration^11^; another study reported drastic mortality in mice treated with antibiotics for gut microbiota depletion and emphasized on the role of gut microbiota-derived single chain fatty acids (SCFAs) in conserving host immune response and offering cardioprotection^24^.

*Fecalibacterium prausnitzii* is a key resident of the gut microbiome, which is known to produce gut protective metabolites^25,26^. Microbial anti-inflammatory molecule (MAM) is one such metabolite that has been identified to exhibit potent anti-inflammatory activity^26^. MAM has also been found to restore intestinal integrity in diabetic mice through the modulation of tight junction transmembrane protein^27^. Our study introduces a novel approach by engineering probiotic EcN to express F. prausnitzii-derived MAM, to ameliorate cardiac translocation of gut bacteria for sustained, *in situ* therapy.

## RESULTS

### 2.1 Natural Mouse Microbiome Translocates to the MI Heart

Our initial objective was to determine whether natural gut bacteria in mice, following systemic leakage, could translocate and colonize the heart after a MI. This inquiry was inspired by cancer research findings, where facultative anaerobes are known to thrive in hypoxic environments like those in solid tumors with hypoxic cores (Supplementary Fig. 1)^15,28–30^. To investigate this, C57BL/6 mice underwent a two-week acclimatization period, followed by coronary artery ligation to induce MI. Based on a previous study indicating peak systemic lipopolysaccharide (LPS) and bacterial leakage within 48-72 hours post-MI, we euthanized the mice 48 hours after surgery^11^. We then conducted bacterial 16S rRNA sequencing on the heart, intestines, and feces to analyze the respective microbiomes. The hearts of sham-operated mice showed negligible bacterial DNA, suggesting minimal to no bacterial presence. In the MI hearts, over 90% of the cardiac microbial composition comprised primarily anaerobic bacteria classes—*Bacteroidia, Clostridia, and Verrucomicrobiae* (Fig. 1A)^31–34^. We observed taxonomic associations between the MI heart, intestines, and fecal microbiomes. There was some correlation between the abundance of bacteria found in the MI hearts and their abundance in the feces and intestines. Notably, despite lower abundance in fecal and intestinal microbiomes, Gammaproteobacteria, which include *E. coli*, were more prevalent in MI hearts (Fig. 1B-C). This aligns with recent findings highlighting differences in intestinal and fecal microbiota^35^. To further explore these associations, we assessed the correlations between the cardiac microbiome and both fecal and intestinal microbiomes. Using Mantel test analyses on unweighted and weighted UniFrac distance matrices, we found moderate, statistically significant correlations between the bacterial populations in MI hearts and intestines (Fig. 1D-E)^36^. A similar pattern of correlation was observed between MI hearts and fecal microbiomes (Fig. 1F-G). To further confirm this phenomenon, we performed a clinically relevant, proof-of-concept experiment using samples from pigs that underwent ischemic reperfusion injury (IRI) surgery (n=3) and naive pigs. PCR for the 16S rRNA gene from DNA extracted from pig hearts identified bacterial DNA in two out of three ischemic hearts 14 days post-surgery, while no bacterial DNA was detected in naïve/healthy hearts (Fig. 1H) (Supplementary Fig. 2).

**Fig. 1.**
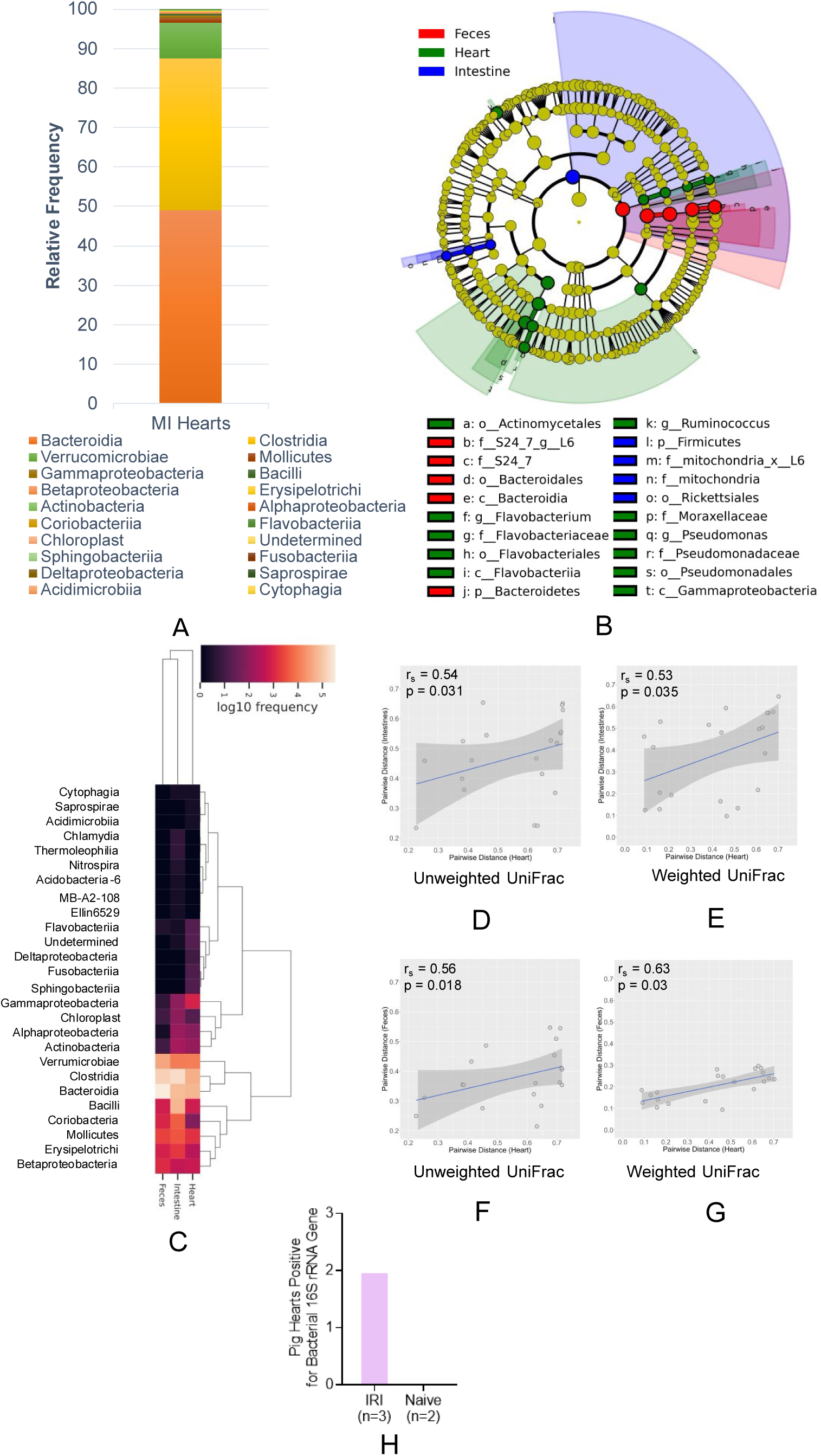
Natural mouse gut microbiome translocates from the gut to the heart after MI. (A) Relative frequency bar chart shows anaerobic classes such as Bacteroidia, Clostridia, and Verrumicrobiaie to be the prominent colonizers of the MI heart, (B) Taxonomic Cladogram from LEfSe showing taxonomic association between MI heart, intestinal, and fecal microbiome samples, (C) Relative abundance of the different microbial classes across MI hearts, intestines, and feces, (D-E) Mantel test analyses of unweighted and weighted UniFrac shows significant correlation in paired samples of MI hearts and intestines (Spearman’s rho (r_s_)=0.54, p=0.031 for unweighted UniFrac; r_s_=0.53, p=0.035 for weighted UniFrac), (F-G) Mantel test analyses of unweighted and weighted UniFrac shows significant correlation in paired samples of MI hearts and feces (r_s_=0.56, p=0.018 for unweighted UniFrac; r_s_=0.63, p=0.03 for weighted UniFrac), (H) PCR for 16S rRNA gene in pig hearts reveals presence of bacteria in ischemic hearts, but not naïve hearts.

### 2.2 EcN Translocates from the Gut to the MI Heart

To further explore the cardiac translocation of gut bacteria post-MI, we employed a tracer bacterium, EcN-Lux, engineered to express luciferase and green fluorescent protein (GFP). This allowed us to track the bacteria using *in vivo* bioluminescence imaging (BLI) as well as for *ex vivo* analysis. After administering EcN-Lux orally to mice, MI surgery was performed. We observed bacterial colonization in the gut as early as 2 days post-administration and continued engraftment beyond 6 days (Fig. 2A). PCR analysis targeting the luxA gene in DNA extracted from the MI hearts revealed the presence of EcN-Lux, unlike in the hearts of sham-operated mice. We noted that cardiac translocation was not uniform across all cases; however, the incidence of EcN-Lux in MI hearts increased over time, reaching 50% by 48 hours (Fig. 2B). Immunohistochemistry against GFP in the ventricular regions of MI hearts confirmed the presence of EcN, validating the translocation phenomenon (Fig. 2C).

**Fig. 2.**
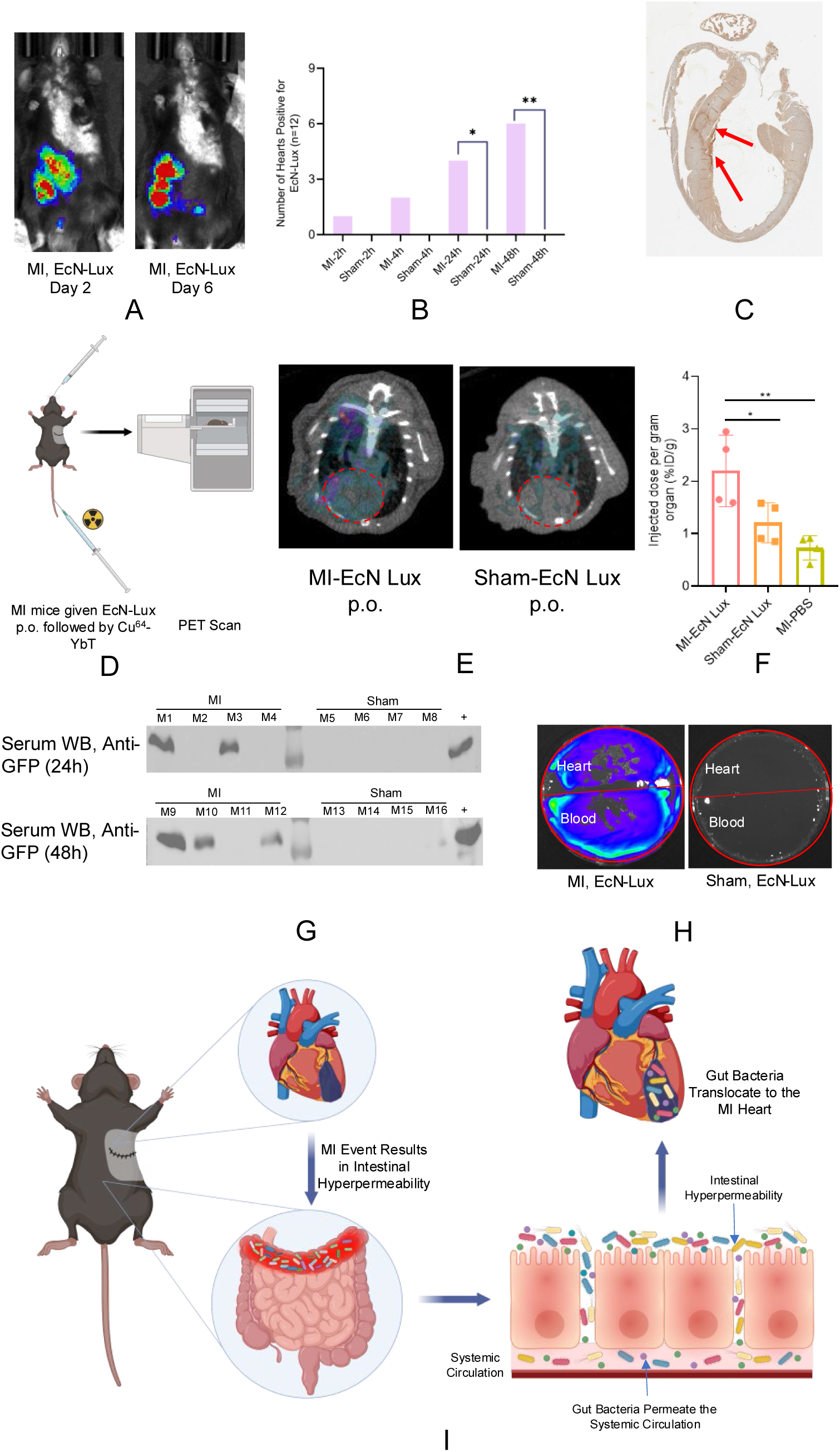
Tracer bacterium EcN-Lux leaks into the systemic circulation and colonizes the heart. (A) Bioluminescence imaging at 2 days and 6 days post administration of EcN-Lux shows continued presence of the administered bacteria in the GI tract, (B) Proportion of heart tissues positive for EcN-Lux increase with time and are consistent with previously published reports that peak systemic leakage of bacteria happens between 48-72h post-MI. Comparison of proportions testing positive for EcN-Lux was done using “N-1” Chi-squared test (*p=0.032, ***p=0.0056), (C) IHC against GFP (Expressed by EcN-Lux) shows cardiac translocation of administered bacteria, (D) Schematic figure representing protocol for siderophore-based PET imaging of live bacteria, (E) ^64^Cu-YbT PET imaging shows higher uptake in the MI heart which confirms presence of live, intact EcN-Lux bacteria. A significantly lower signal was detected in the sham hearts, (F) Biodistribution studies for ^64^Cu-YbT confirms that MI hearts have a significantly higher presence of ^64^Cu-YbT complex indicating presence of live, intact bacteria in the MI heart. Data represented as mean ± SD (n=4) and compared using one-way ANOVA (Tukey’s multiple comparison test) (**p=0.0043, *p=0.0373), (G) Immunoblotting of serum of MI and sham mice given oral EcN-Lux shows presence of GFP and confirms leakage of bacteria into the systemic circulation (H) Culturable EcN-Lux was found in both the blood and the hearts of MI mice given EcN-Lux orally. IVIS imaging of heart extract and blood culture plates confirmed presence of EcN-Lux in hearts and blood of MI mice but not in shams, (I) Schematic illustrating the leakage and translocation of gut bacteria to the MI heart.

To assess whether the translocated bacteria were indeed live bacteria, we employed siderophore-based PET imaging and radionuclide-based biodistribution studies. Siderophores are small metal-chelating molecules secreted by bacteria to uptake iron and other nutrient metals^37,38^. We have previously developed a siderophore-based radio-imaging technique using positron emission tomography/computed tomography (PET/CT) allowing us to non-invasively image live bacteria with high specificity and species-selectivity in the body^39^. A complex of radioactive copper, copper-64 (^64^Cu) and the siderophore, yersiniabactin, was administered 24h post-surgery (Fig. 2D). PET imaging showed a significantly higher presence of ^64^Cu-Yersiniabactin (^64^Cu-YbT) in the MI heart compared to sham, indicating the presence of live EcN (Fig. 2E). *Ex vivo* biodistribution studies were also performed to assess residual radioactivity. MI hearts were found to have a significantly higher ^64^Cu-YbT per gram of tissue compared to sham hearts which further corroborates presence of live EcN in the heart (Fig. 2F)^39^.

Furthermore, immunoblotting of serum from MI mice administered with EcN-Lux at 24h and 48h post-surgery revealed presence of bacterial GFP in the serum of the MI mice suggesting leakage of EcN-Lux into the systemic circulation (Fig. 2G). Additionally, plating of heart and blood extracts from MI mice on ampicillin-selective media resulted in bacterial growth, as evidenced by bioluminescence, while sham mice showed no such growth (Fig. 2H). These findings collectively support the hypothesis that gut bacteria, following MI, can indeed translocate and colonize the heart (Fig. 2I).

### 2.3 Systemically Administered EcN Colonize and Proliferate in the MI Heart

We further validated the propensity of anaerobic bacteria like EcN to localize in ischemic cardiac tissue. To mimic gut bacteria leakage, we administered EcN-Lux intravenously right after inducing MI in mice (Fig. 3A). BLI distinctly showed the localization and colonization of EcN-Lux in the MI hearts as early as 24 hours post-MI. In contrast, sham-operated mice that received intravenous EcN-Lux displayed no bioluminescent signals (Fig. 3B). This finding underscores the ability of EcN, a bacterium naturally found in the human gut, to specifically colonize and proliferate in the infarcted heart. To reinforce these observations, we conducted ^64^Cu-YbT PET imaging, for imaging live bacteria. The PET imaging and subsequent radioactivity-based biodistribution studies revealed a significantly higher accumulation of the ^64^Cu-YbT probe in the hearts of MI mice compared to the sham group (Fig. 3C-D). This indicates the presence of live EcN-Lux in the MI hearts. Furthermore, IHC analysis of the infarcted heart tissues demonstrated a clear presence of the bacteria. This finding aligns with the BLI and PET imaging results and solidifies the conclusion that EcN can achieve targeted colonization specifically in the hearts affected by MI (Fig. 3E).

**Fig. 3.**
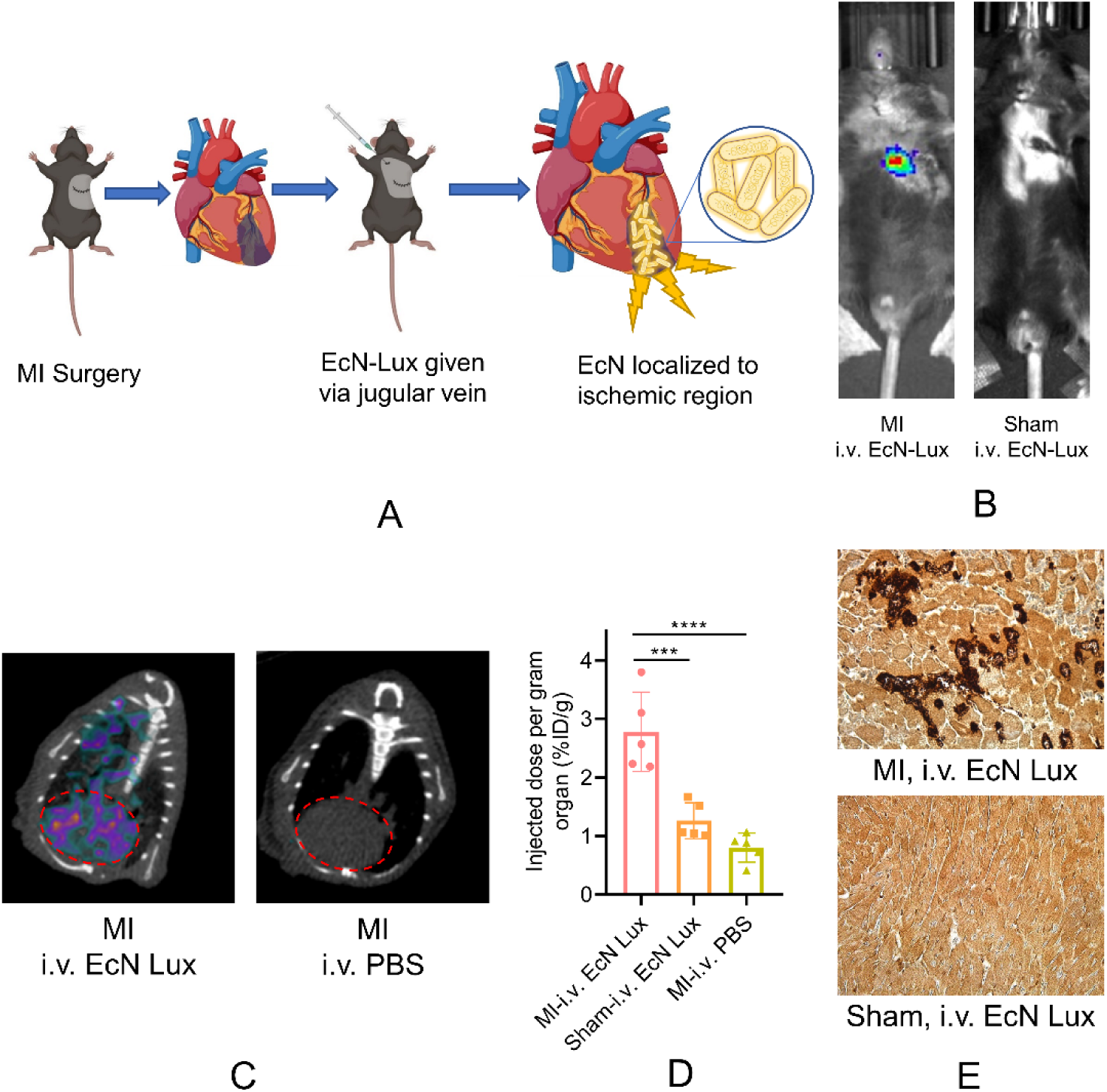
Simulation of gut bacteria leakage by intravenous administration of EcN-Lux shows clear localization of bacteria in the heart tissue. (A) Schematic figure of the method used to simulate gut leakage of live bacteria, (B) Bioluminescence imaging shows that i.v. administration of EcN-Lux results in its colonization and proliferation in MI hearts but not shams, (C) ^64^Cu-YbT PET imaging in MI mice administered intravenous EcN-Lux confirms colonization of the MI heart by live, intact bacteria in the MI heart (Left). No signal was observed in mice administered PBS (Right), (D) Biodistribution studies using ^64^Cu-YbT revealed higher radioactive signal in hearts of MI mice given intravenous EcN-Lux compared to controls. Data represented as mean ± SD (n=4) and compared using one-way ANOVA (Tukey’s multiple comparison test) (***p=0.0005, ****p<0.0001), (E) IHC against GFP on heart tissue of MI mice given i.v. EcN-Lux shows clear presence of GFP in the MI heart but not in sham.

### 2.4 Recombinant EcN produces MAM protein

Drawing from our findings and existing research, we developed a novel microbiome-based construct designed to prevent the leakage of gut microbiota into the systemic circulation^11,12,40^. We engineered EcN to produce MAM protein derived from *F. prausnitzii*^25,27^. The constructed pBbA19a plasmid containing 6xHis tagged MAM protein was transformed into EcN using heat shock and transformants were obtained using ampicillin selection (Fig 4A). Sanger sequencing of transformed colonies confirmed successful incorporation of pBbA19a-MAM plasmid into EcN. Immunoblotting of EcN-MAM was performed to validate the synthesis of MAM protein by the bacteria. SDS-PAGE of bacterial protein followed by immunoblotting using anti-6xHis Tag antibody demonstrated presence of MAM in the bacterial protein extract (Fig. 4B).

**Fig 4.**
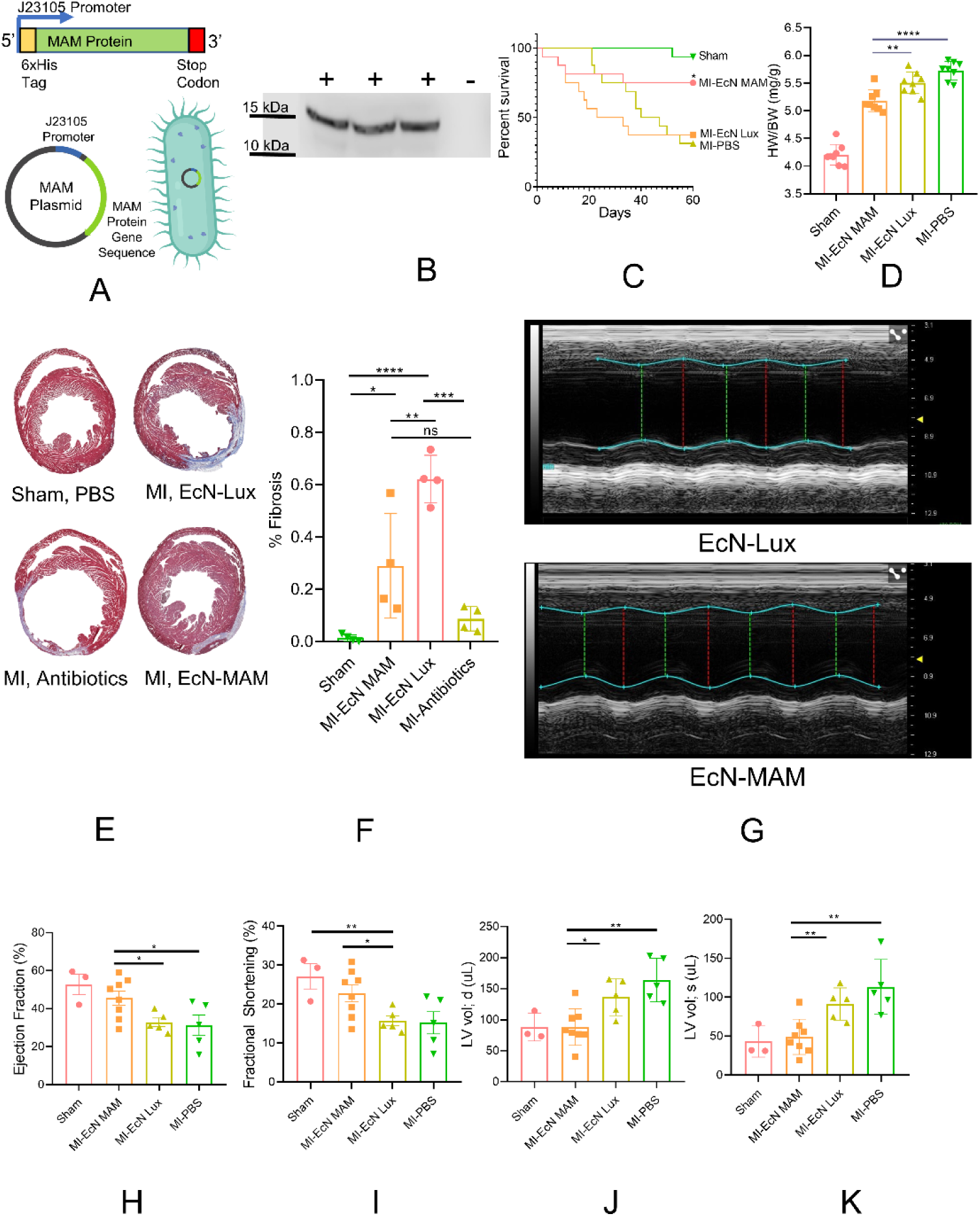
EcN-MAM results in increased survival and better cardiac outcomes in MI mice. (A) Schematic of the gene circuit, plasmid map, and engineered EcN-MAM, (B) Immunoblotting of bacterial protein extract demonstrates that the engineered EcN-MAM construct produces the Fecalibacterium prausnitzii-derived MAM protein, (C) Kaplan-Meier survival curve showing significantly longer survival in MI mice treated with EcN-MAM vs. controls. Groups were compared using logrank (Mantel-Cox) test (n=12, *p=0.0492 for MI-EcN MAM vs. MI-EcN Lux, *p=0.043 for MI-EcN MAM vs. MI-PBS), (D) Mouse heart weight to body weight ratio (HWBR) is significantly reduced in mice treated with EcN MAM vs. mice treated with EcN Lux indicating lower fibrosis. Data represented as mean ± SD (n=8) and compared using one-way ANOVA (Tukey’s multiple comparison test) (**p=0.0085, ****p<0.0001), (E-F) Masson’s trichrome staining reveals significantly lower cardiac fibrosis in mice treated with EcN MAM compared to EcN Lux controls. Data represented as mean ± SD (n=4) and compared using one-way ANOVA (Tukey’s multiple comparison test) (**p=0.0061, ***p=0.0001, ****p<0.0001), (G-K) Echocardiographic images and assessment of cardiac function at 4 weeks post-surgery shows significantly better cardiac parameters in EcN MAM treated mice. Data represented as mean ± SEM and compared using unpaired t-test (*p<0.05, **p<0.01).

### 2.5 Therapy with EcN-MAM Results in Increased Survival, Reduced Post-MI Cardiac Fibrosis, and Better Cardiac Function

We conducted *in vivo* experiments to assess the effectiveness of EcN-MAM’s efficacy in the mouse model of MI. We treated mice with either EcN-MAM or EcN-Lux prior to inducing MI, then tracked their survival over a 60-day period post-surgery. Our findings revealed a significant improvement in survival rates for the EcN-MAM group compared to those treated with EcN-Lux or a PBS control. Specifically, over 50% of the EcN-MAM treated mice survived beyond the 60-day mark, leading to an undefined median survival time, whereas the median survival times for the EcN-Lux and PBS groups were 28 and 43.5 days, respectively (Fig. 4C).

In a separate group of mice, we evaluated heart health 28 days after surgery by measuring the heart weight to body weight ratio (HWBWR). Mice treated with EcN-MAM showed a significantly lower HWBWR, indicating reduced heart hypertrophy and better cardiac healing compared to those treated with EcN-Lux (Fig. 4D). We also assessed cardiac fibrosis using Masson’s trichrome staining, a standard method for this purpose. To understand the role of gut bacteria-mediated inflammation in cardiac healing, we included an additional group of MI mice pre-treated with broad-spectrum antibiotics to deplete their gut microbiome. The results showed significantly less fibrotic tissue in the hearts of EcN-MAM treated mice compared to those treated with EcN-Lux. Interestingly, mice with a depleted gut microbiome also exhibited reduced cardiac fibrosis,^11^ although this difference was not statistically significant when compared to the EcN-MAM group (Fig. 4E-F). This suggests that EcN-MAM treatment effectively reduces post-MI cardiac fibrosis and hypertrophy. Further, we used echocardiography to evaluate cardiac function in the mice treated with EcN-MAM. The echocardiographic analysis showed that the EcN-MAM group had significantly better cardiac function across various parameters, including ejection fraction, fractional shortening, and left ventricular volume during diastole and systole, compared to both the untreated and EcN-Lux treated mice (Fig. 4G-K).

### 2.6 EcN-MAM Mitigates Inflammation in the Infarcted Heart

We analyzed immune cell dynamics in the hearts of EcN MAM-treated mice at 72h post-MI. The Uniform Manifold Approximation and Projection (UMAP) images (Fig. 5A) displayed a noticeable shift in the balance of immune cells compared to naïve controls, with an expected increase in monocytes and neutrophils in the MI-affected groups. Further detailed analysis revealed that the EcN-MAM group exhibited significantly lower numbers of monocytes, macrophages, neutrophils, and particularly activated neutrophils (Siglec-F+) when calculated as a percentage of total live cells (Fig. 5B–5F). The presence of Siglec-F+ neutrophils is notable since they are known to exacerbate renal fibrosis by promoting collagen 1 expression, higher levels of profibrotic inflammatory cytokines, and increased hypersegmentation^41^. This suggests that treatment with EcN-MAM results in a markedly reduced proinflammatory response, likely due to decreased systemic bacterial permeation and the subsequent immune response.

**Fig. 5.**
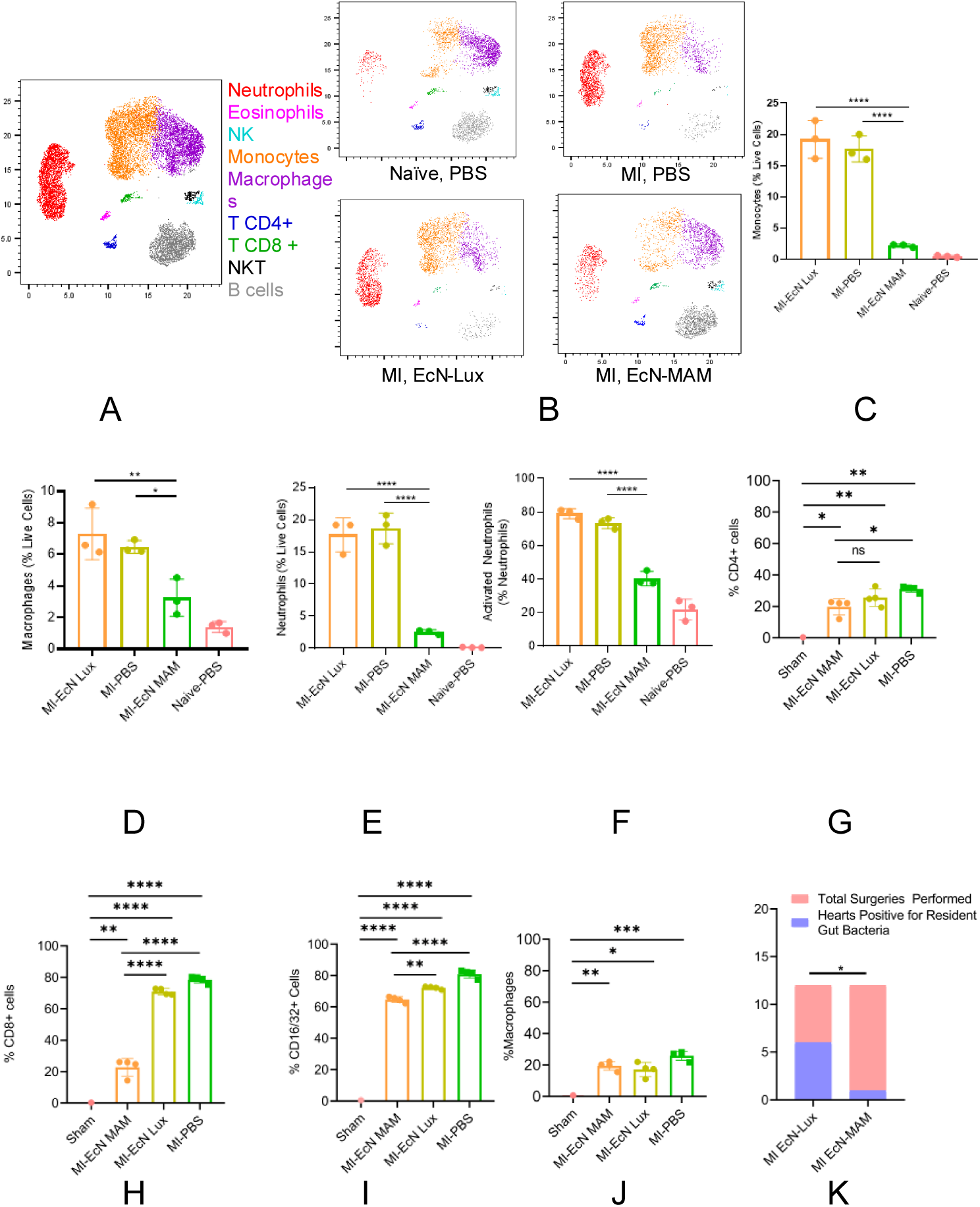
EcN-MAM treated MI mice result in better cardiac immune response 48h post-surgery. (A) UMAP representation of different treatment and procedure groups (Expressed in CD45+ immune cells) indicates massive recruitment of monocytes and neutrophils in MI group, (B-F) Flow cytometric analysis shows EcN MAM treated mice have significantly fewer numbers of monocytes (C) and macrophages (D) at 72h timepoint. MI mice treated with EcN MAM also had significantly lower numbers of neutrophils (E) and activated neutrophils (F) compared to EcN Lux and PBS treated controls. Data represented as mean ± SD (n=3) and compared using one-way ANOVA (Tukey’s multiple comparison test) (*p<0.05, **p<0.01, ***p<0.001, ****p<0.0001), (G-J) Flow cytometric analyses of mouse spleens indicating changes in cell populations of CD4+ cells (G), CD8+ cells (H), CD16/32+ cells (I), Macrophages (J), (K) EcN-MAM was found in a significantly lower proportion of MI hearts compared to EcN-MAM in mice treated with respective bacteria. Comparison of proportions testing positive for respective gut bacteria was done using “N-1” Chi-squared test (*p=0.0279)

We further investigated changes in immune population in mouse spleens after MI. We observed significant upregulation of both CD4+ and CD8+ cells after MI, indicating recruitment of both helper and cytotoxic T-cells. However, we observed both populations of T-cells to be less upregulated in the EcN-MAM group compared to EcN-Lux and PBS groups (Fig. 5G–5H). A similar trend was observed in the case of CD16/32+ cells, which comprise of macrophages, monocytes, neutrophils, and NK cells^42,43^. We observed CD16/32+ cell population to be significantly lower in the EcN-MAM group compared to the EcN-Lux and PBS groups (Fig. 5I). However, no significant difference was observed in macrophage populations in the spleen of the MI mice (Fig. 5J). Overall, investigation into the immune cell dynamics in the spleen revealed a systemic dampening of the immune response, in the EcN-MAM group, compared to the EcN-Lux and PBS groups, which aligns with reduced inflammation in cardiac tissue. These observations correlate with the immune cell population observed in the heart, indicating a less intense proinflammatory physiological response post-MI, reducing the risk of maladaptive cardiac remodeling^44,45^.

To investigate whether the therapeutic effect of EcN-MAM was a result of the engineered bacteria translocating to the heart or locally releasing MAM protein in the gut, we conducted PCR studies. These studies revealed that a significantly lower number of MI hearts tested positive for EcN-MAM compared to EcN-Lux, suggesting that the primary action of EcN-MAM occurs at the intestinal level, in line with previous research (Fig. 5K)^25,27^. This led us to pivot our focus towards examining the impact of EcN-MAM at the intestinal level.

### 2.7 EcN-MAM Restores the Post-MI Gut Barrier

As previously discussed, MI significantly compromises gut barrier integrity, leading to intestinal hyperpermeability. MAM protein has previously been shown to restore the gut barrier after a non-MI related pathological insult, such as colitis^25,27^. We analyzed the expression of key intestinal tight junction proteins, occludin and zona occludens-1 (ZO-1), in large intestinal tissue samples collected 48 hours after surgery. Our results showed that mice treated with EcN-MAM exhibited enhanced gut barrier integrity, as evidenced by higher levels of occludin and ZO-1, compared to those treated with EcN-Lux and untreated mice. This indicates a potential protective role of EcN-MAM in preserving the gut barrier (Fig. 6A). Given that intestinal hyperpermeability can lead to the leakage of bacterial products like lipopolysaccharides (LPS) into the bloodstream, triggering endotoxemia and inflammation,^46,47^ we measured the levels of Lipopolysaccharide Binding Protein (LBP) in the serum of mice. LBP, a response marker to LPS, plays a vital role in innate immune response to inflammation^48^. Our findings revealed that mice treated with EcN-MAM had significantly lower serum LBP levels compared to the EcN-Lux group, underscoring the protective effect of EcN-MAM. Interestingly, the LBP levels in MI mice treated with EcN-MAM were comparable to those in sham mice (Fig. 6B). We also measured C-reactive protein (CRP) levels, a well-established marker of systemic inflammation^49,50^. Although murine CRP is less sensitive than its human counterpart, it still serves as a reliable indicator of post-MI inflammatory response^51,52^. Mice treated with EcN-MAM showed significantly lower CRP levels compared to those treated with EcN-Lux. Furthermore, treatment with broad-spectrum antibiotics also resulted in lower CRP levels, supporting the hypothesis of gut bacteria-mediated inflammation (Fig. 6C). Finally, we conducted biodistribution studies to address safety concerns about EcN-MAM potentially translocating to other major organs. These studies confirmed that EcN-MAM did not translocate to any major organs (Supplementary Fig. 3), suggesting its targeted action in the gut. Therefore, our results lead us to conclude that EcN-MAM primarily functions by maintaining or restoring the integrity of the intestinal barrier post-MI (Fig. 6D).

**Fig. 6.**
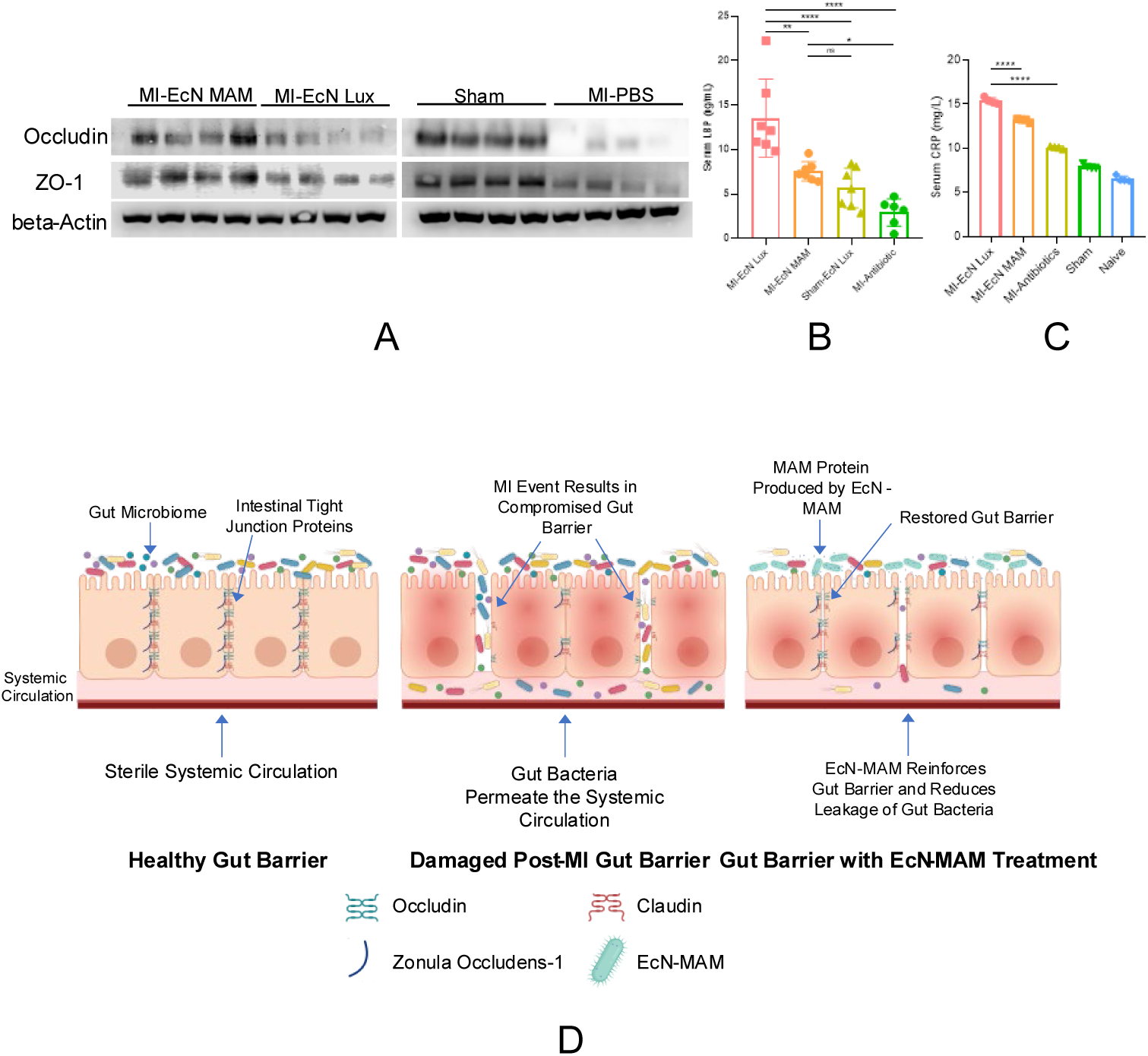
EcN-MAM treatment protects against disruption of the gut barrier post-MI. (A) Immunoblotting of occludin and ZO-1 shows MI mice treated with EcN MAM have higher levels of intestinal tight junction proteins vs. MI mice given EcN Lux or PBS. Loading control (beta-actin) was run on a duplicate gel, (B) MI mice given EcN MAM have significantly lower levels of serum LBP indicating less leakage of gut bacteria into the systemic circulation. Data represented as mean ± SD (n=4) and compared using one-way ANOVA (Tukey’s multiple comparison test) (*p=0.0242, **p=0.0019, ****p<0.0001), (C) Serum C-Reactive Protein (CRP) levels are significantly elevated in control mice given EcN Lux, but are significantly resolved in mice given EcN-MAM or mice with antibiotic-induced intestinal microbiome depletion. Data represented as mean ± SD (n=4) and compared using one-way ANOVA (Tukey’s multiple comparison test) (****p<0.0001), (D) Schematic illustrating the proposed mechanism of action of EcN-MAM

## DISCUSSION

Myocardial infarction is a severe cardiac event that results in acute global changes in the body. Ischemia results in necrosis of myocardial tissue which ultimately causes cardiac dysfunction. Association between chronic heart failure (CHF) and intestinal dysfunction was discovered over a decade ago when CHF patients were found to have a 35% and 210% increase in small and large intestine permeability respectively compared to control patients^53^. Since then, numerous clinical and pre-clinical studies have elucidated the MI heart-gut axis as well as the role of the gut microbiome in the pathology of MI^12,54^. Our 16S rRNA analysis revealed presence of bacteria belonging to the natural murine microbiome in the hearts of MI mice. Consistent with our hypothesis, bacteria belonging to anaerobic classes such *as Bacteroidia, Clostridia*, and *Verrucomicrobiae* were highly abundant in the MI hearts. Bacteria belonging to *Bacteroidia* and *Clostridia*, which are present in significantly greater abundance in the large intestines, further indicated that large intestine is the primary region affected by post-MI intestinal hyperpermeability consistent with clinical reports^53^. Microbiome research has revealed both inter-subject and intra-subject statistical differences in the fecal and intestinal microbiomes^55,56^. Furthermore, fecal samples have been found to not represent the entire gut microbiome^55^.

Therefore, we applied Mantel test analyses on both unweighted and weighted UniFrac distance matrices. The unweighted UniFrac provides insights into the presence or absence of bacterial species, highlighting low abundance features, while the weighted version considers the relative abundance of species/taxa^57^. Mantel test allows us to incorporate paired analyses between intra-subject microbiomes. Our findings showed significant correlations between the microbiomes of the MI heart and intestines, as well as the MI heart and fecal microbiomes. This correlation uncovers a new aspect of MI pathophysiology, confirming the translocation and colonization of gut bacteria in the MI heart.

To further substantiate our hypothesis, we employed EcN-Lux, a tracer bacterium capable of producing bioluminescence and GFP. Our studies indicated a time-dependent correlation in the cardiac presence of bacteria post-surgery. This aligns with previous findings that intestinal permeability and systemic LPS levels peak within 48-72 hours post-MI^11^. We have previously established a method for detecting live, intact bacterial cells in the body using PET imaging^39^. Using PET imaging with a ^64^Cu-YbT complex, we tracked live bacterial translocation. The uptake of this complex by live bacteria, making them detectable via PET imaging, confirmed the colonization of the MI heart by live bacteria.

*E. coli*, particularly the facultative anaerobe EcN, thrives in both aerobic and anaerobic environments and is a pioneer colonizer of the human intestines, present in over 90% of the population^58^. Previous studies have shown that EcN can colonize hypoxic regions like solid tumors when administered intravenously^15,59^. Thus, we postulated that the ischemic and hypoxic conditions of the injured myocardium provide an ideal environment for EcN. Our experiments with intravenously administered EcN-Lux, leading to a distinct bioluminescent signal in the cardiac region of MI mice but not shams, further validated this theory.

Consequently, we developed a probiotic bacterial construct to mitigate systemic bacterial permeation. Gut-derived low-grade endotoxemia is a critical concern in cardiovascular diseases, exacerbating conditions like atherosclerosis and thrombosis^46^. Clinical studies have shown elevated LPS in coronary thrombi of ST-elevation myocardial infarction (STEMI) patients compared to stable angina patients,^60^ and its presence in the bloodstream has been linked to major adverse cardiovascular events (MACE) in atrial fibrillation patients^61^. This LPS influx is largely attributed to increased intestinal permeability. Hence, we focused on targeting intestinal hyperpermeability with our construct.

MAM is a small 15 kDa protein which was identified to be secreted by *Fecalibacterium prausnitzii*. F. prausnitzii is a strictly anaerobic bacteria, the intestinal population of which is inversely correlated with gut disorders. MAM has been found to be one of the prominent metabolites behind F. prausnitzii’s gut protective actions and has demonstrated to be effective in treating colitis as well as restoring intestinal integrity in different mouse models^26,27^. We engineered EcN to produce MAM, hypothesizing that this would reduce post-MI bacterial systemic permeation by protecting the gut barrier. The use of a live biotherapeutic like EcN offers the advantage of sustained, *in situ* therapeutic production.

Our results showed that MI mice treated with EcN-MAM had significantly improved survival, reduced heart to body weight ratio, less cardiac fibrosis, and better echocardiographic parameters than those treated with EcN-Lux. To unravel the therapeutic mechanism, we considered the acute pro-inflammatory response following MI. This response, involving neutrophils, cytokines, inflammasomes, and toll-like receptor activation, typically lasts about three days and transitions into an anti-inflammatory reparative phase. Imbalances in this transition can lead to prolonged inflammation, resulting in fibrosis and poorer outcomes^62^. We observed reduced activation of proinflammatory cells in EcN-MAM treated MI mice in both heart and spleen, aligning with studies linking early inflammatory biomarkers to increased mortality^63,64,65^. Despite its engineered nature, EcN-MAM does not elicit a robust proinflammatory immune response, either locally in the gut or systemically. The observed mild immune response, particularly in T-cell (CD4+ and CD8+) and CD16/32+ cell populations, suggests that the engineered strain is well-tolerated by the host immune system.

Initially suspecting that EcN-MAM’s action might be due to heart colonization, we found no presence of EcN-MAM in the MI heart, leading us to conclude that the therapeutic benefit stemmed from diminished immune activation due to reduced systemic bacterial permeation. Immunoblotting showed enhanced gut integrity in EcN-MAM-treated mice, indicated by higher levels of occludin and ZO-1. Elevated serum LPS levels, a common feature in cardiovascular disease patients, trigger TLR4-mediated inflammatory signaling, detrimental to cardiomyocytes and associated with MACE^61,66^. This endotoxemia results in chronic systemic inflammation, cardiac fibrosis, and heart failure. A clinical study with 2568 participants revealed a significant association between high serum lipopolysaccharide binding protein (LBP) levels and the development of cardiovascular disease^67^. We performed ELISA to assess the levels of systemic LBP in MI mice as a surrogate and more reliable marker for systemic LPS. We observed that EcN-MAM treated mice had significantly lower levels of serum LBP compared to mice given control EcN. We also found that depletion of the intestinal microbiome through administration of antibiotic cocktails resulted in a very significant decline in serum LBP levels corroborating the role of gut bacteria as a cause for elevated LBP numbers. We also performed an ELISA for CRP to investigate the state of systemic inflammation in the MI mice. CRP, an acute phase reactant protein and a gold standard of inflammation in the body, has also been found to predict in-hospital outcome and long-term mortality. A clinical study by Mani et al. with 5145 patients revealed significant and independent associations between high sensitivity CRP (hsCRP) and major adverse cardiac event (MACE), cardiovascular death, and all-cause death^68–70^. We found that mice treated with EcN-MAM had significantly lower levels of serum CRP compared to control mice and antibiotic cocktail mediated microbial depletion demonstrated a further decrease in serum CRP. Depletion of the gut microbiome has been investigated as a strategy to mitigate gut bacteria-mediated post-MI systemic inflammation, however, as discussed previously, this approach has yielded mixed results^11,71^. Additionally, depletion of the gut microbiome using a cocktail of broad-spectrum antibiotics is not a viable clinical strategy due to risk of antibiotic resistance, superinfections, potential drug-drug interactions and increased side effects.

To our knowledge, our study is the first to discover the phenomenon of gut translocating and colonizing the infarcted heart. We are also the first to report the use of a recombinant probiotic platform secreting a defined therapeutic protein, MAM, to target intestinal hyperpermeability and protect against excessive systemic permeation of gut microbiota. However, there are several shortcomings to this study. Future studies require further evaluation on the role of bacteria translocated to the heart in the pathophysiology and recovery of myocardial infarction.

Additionally, evaluating whether a certain portion of therapeutic bacteria that translocated to the heart can likely exert a direct therapeutic effect on the cardiac tissue would be helpful in optimizing therapeutic benefit and designing future live biotherapeutic products. Evaluating the effect of EcN-MAM in an ischemia-reperfusion injury model will be beneficial due to the distinct nature of injury in this model, compared to the total occlusion model. Although we used EcN – a known probiotic – as the vector, EcN itself produces LPS and therefore constructs made from other probiotic bacteria such as *Lactococci* or *Bacilli* can also be evaluated. In the current study, we utilized male mice and therefore, studies need to be repeated in female mice to evaluate if sex differences result in any changes with respect to cardiac translocation of gut bacteria or efficacy of EcN-MAM.

In conclusion, our research demonstrates that MI not only leads to systemic circulation of live bacteria but also their colonization in injured cardiac tissue. Additionally, oral administration of EcN-MAM significantly reduces post-MI cardiac inflammation and fibrosis.

## METHODS

### IACUC Statement

Animals were maintained in an Association for the Assessment and Accreditation of Laboratory Animal Care approved facility (Assurance # D16-00190) in accordance with current regulations of the U.S. Department of Agriculture and Department of Health and Human Services. Experimental methods were approved by and in accordance with institutional guidelines established by the Institutional Animal Care and Use Committee (approved protocol numbers: 23-03-30-02).

Experiments in pigs were performed using protocols that adhered to guidelines and approved by Institutional Animal Care and Use Committee at the University of Arizona and to 2019 NIH guidelines for care and use of laboratory animals^72^.

### Myocardial Infarction Procedure

Eight to ten weeks old male C57/BL6 mice were shaved around the thoracic and neck regions. Sterile technique was followed per institutional protocols. Mice were pre-emptively given 1mg/kg Buprenorphine Extended Release (Wedgewood Pharmacy) subcutaneously, then every 48-72 hours post-operatively as needed. Mice were anesthetized with 2-2.5% isoflurane (inhale to effect) during the procedure. The animals were intubated and placed on a ventilator (MiniVent Type 845, Hugo Sachs Elektronik, March-Hugstetten, Germany). Body temperature was maintained at 37 °C. A thoracotomy was performed at the 4th intercostal space in order to visualize the LAD. For occlusion, an 8/0 nylon suture was placed around the LAD approximately 1mm distal to the left atrial appendage. Occlusion was confirmed by blanching of the myocardium and a ST elevation indicated on the ECG (Supplementary Fig. 4). The chest wall was closed with absorbable suture and the skin with non-absorbable suture. Upon spontaneous respiration, the mice were extubated and moved to a temperature-controlled area with oxygen supplementation until they regained consciousness. The sham surgery for this model involved all the above steps including passing the suture around the LAD, however the artery was not ligated.

### Ischemia Reperfusion Injury Model in Pigs

Ischemia reperfusion injury was induced in pigs as per protocol described by Skaria et al^72^.

### DNA Extraction and Polymerase Chain Reaction (PCR)

Mouse hearts were extracted aseptically after perfusion with cold sterile PBS and DNA extraction was carried out using the DNEasy® Blood & Tissue Kit (Qiagen, Germantown, MD) as per the manufacturer’s instructions. PCR was performed using DreamTaq® DNA polymerase (ThermoScientific^TM^, Waltham, MA) for a 500 bp fragment of the luxA gene with primers from Integrated DNA Technologies, Coralville, IA. Gel electrophoresis of the PCR product was performed on a 1.5% agarose gel.

For microbiome analysis, each mouse was placed in a sterile cage and their feces were collected. Mice were subsequently euthanized, and organs were dissected using the aforementioned protocol. Heart and intestinal DNA was extracted using Molzym Ultra-Deep Microbiome Prep Kit (Molzym GmbH & Co. KG, Bremen, Germany). Fecal DNA was extracted using QIAamp PowerFecal Pro DNA Kit (Qiagen, Germantown, MD). The extracted DNA was submitted to the Cincinnati Children’s Hospital Medical Center DNA Core facility for 16S metagenomics sequencing. Briefly, quality control of DNA samples was performed using 515F-806R primers. The V4 hypervariable region of the 16S rRNA gene was amplified and sequenced as per Illumina MiSeq’s 250PE run protocol. The generated data was analyzed using standard QIIME2 protocols^73^. UniFrac figures and Mantel test analyses was done in RStudio using the vegan package.

### Bacterial engineering

Recombinant *E. coli* Nissle 1917 (EcN) producing GFP, and luciferase was engineered using the pAKgfplux1 plasmid (Gift from Attila Karsi, Addgene plasmid #14083).

*E. coli* Nissle 1917 (EcN) was engineered to produce *Fecalibacterium prausnitzii*-derived microbial anti-inflammatory protein (MAM)^26^. Double-stranded DNA fragment encoding for MAM protein was obtained from Integrated DNA Technologies, Coralville, IA. A 6xHis tag was added to the DNA sequence to aid in downstream studies. Fragments were amplified using PCR and ligated into a pBbA19a backbone^74^. Chemically competent EcN were transformed using heat shock and transformed colonies were grown and their sequence was corroborated using Sanger sequencing.

### EcN Gut Colonization Model

Mouse drinking water was supplemented with 0.25 mg/mL ampicillin. Mice were given 10^8^ colony forming units (cfu) of log-phase *E. coli* Nissle 1917 through oral gavage. Bacteria were given one dose every week.

Due to the short gut transit time in mice, competition from existing mouse microbiota, and egestion of administered bacteria prior to achieving gut colonization, we were unable to achieve uniform colonization without frequent readministrations. We evaluated supplementation of mouse water with 0.25 mg/mL ampicillin prior to bacterial administration. We observed that bacterial administration one day after supplementing the water with ampicillin with continuous supplementation post-administration allows for successful gut colonization of our tracer bacterium EcN-Lux (*E. coli* Nissle 1917 expressing pAkgfplux1 plasmid). Since EcN-Lux bears resistance to ampicillin, ampicillin administration not only ensures the depletion of some gut microbes which provides GI tract area for EcN-Lux to colonize, but also ensures that our administered bacterium does not lose its plasmid after several generations of multiplication. We observed that EcN-Lux was able to maintain successful gut colonization up to one week after oral administration of EcN with ampicillin supplementation of water. *Ex vivo* bioluminescence imaging (BLI) of the GI tract indicated colonization of the distal small intestine and colon by EcN-Lux – the two regions of the GI tract which have significantly increased permeability post-MI (Supplementary Fig. 5).

### Microbiome Depletion

Gut microbiome depletion was carried out as per previously described method^75^. Briefly, mice were given drinking water supplemented with 1 mg/mL ampicillin and 0.575 mg/mL enrofloxacin for a period of 8 to ensure depletion of the gut microbiome.

### Histopathology

After euthanasia via carbon dioxide inhalation, mouse hearts were isolated, weighed, and fixed in neutral buffered formalin (NBF) and embedded in paraffin. 6-micron sections of the tissue were cut by Cincinnati Children’s Hospital Medical Center Department of Pathology Research Core (Cincinnati, OH, USA). Immunohistochemistry was performed using anti-GFP antibody (Abcam).

Transverse sections of MI hearts were stained for Masson’s trichome to assess post-MI fibrosis. Six sections were cut per heart. Trichrome staining was quantified as per protocol described by Kennedy et al.^76^

### Immunoblotting to confirm bacterial synthesis of MAM protein

Overnight culture of EcN-MAM was inoculated in 100 mL of terrific broth (1:100 dilution) and incubated at 37 °C at 200 rpm in a shaker-incubator for 3 hours. Subsequently, the culture was centrifuged at 3000g for 15 min at 4 °C, followed by careful removal of the supernatant. To allow for efficient protein extraction, the bacterial pellet was frozen in dry ice for 1h and then allowed to thaw on ice. Bacterial protein was extracted using Bacterial Protein Extraction Reagent (B-PER^TM^) supplemented with DNase I, lysozyme, and Halt^TM^ protease inhibitor cocktail as per the manufacturer’s instructions. The extracted protein was mixed with Tricine Sample Buffer (BioRad, Hercules, CA) as per manufacturer’s instructions and denatured at 95 °C for 10 min.

The protein samples were loaded into a 16.5% tris-tricine gel and SDS-PAGE was performed in a tris/tricine/SDS buffer system (Biorad, Hercules, CA). The gel was transferred onto a 0.2 µm nitrocellulose membrane and blocked using 5% bovine serum albumin (BSA) in tris-buffered saline (TBS). The membrane was then incubated with HRP conjugated 6xHis-tag monoclonal antibody (Proteintech, Rosemont, IL) at 1:10,000 dilution in TBS-Tween. Membrane was washed three times with TBS-Tween followed by addition of Supersignal^TM^ West Pico PLUS Chemiluminescent Substrate (Thermo Scientific, Waltham, MA). The blot was detected using the C-DiGit Blot Scanner (LI-COR Biosciences, Lincoln, NE).

### Echocardiography

Cardiac function was assessed via echocardiography using a Vevo 2100 imaging system (VisualSonics, Toronto, Canada) as previously described^77^. Briefly, mice were anesthetized using isoflurane inhalation, chest hair was removed using a depilatory cream, and each mouse was subsequently placed on a heated platform maintained at 38 °C with paws in contact with the electrocardiography leads. Mouse core body temperature was monitored using a rectal probe and maintained at 36-37 °C using a heat lamp and the heated platform. M-mode echocardiography was performed at the parasternal short axis and images were analyzed for various parameters of cardiac function such as cardiac output, fractional shortening, ejection fraction, and LV mass. Mice were imaged before and after MI surgery followed by weekly electrocardiography imaging. Post euthanasia, mouse heart weights were determined for normalization and assessment of fibrosis and hypertrophy^78^.

### Flow Cytometry

Heart Tissue: Perfused and harvested hearts were minced into 1mm pieces using a razor blade. The minced hearts were washed three times with phosphate-buffered saline (PBS) to remove intravascular cells. The hearts were then digested for 45 minutes at 37 °C under agitation, in Hank’s Balanced Salt Solution (HBSS) solution containing 1mg/ml Collagenase IV (Gibco) and 10ug/ml DNase I (Sigma-Aldrich). Cell suspensions were filtered through a 70µm strainer, and debris were removed using Debris removal solution (Miltenyi), following the supplier’s protocol. For immunophenotyping by flow cytometry, two million cells per sample were used. The single cell suspensions were first labeled for 30 minutes at 4 °C with LIVE/DEAD Fixable Blue (Invitrogen), to separate live cells from dead cells during analysis. After two washes, the samples were incubated for 30 minutes at 4 °C with the following antibodies: anti-CD45-BUV496 (BD Bioscience, 30-F11), Anti-CD3-APC (Biolegend, 17A2), anti-CD11b-PacificBlue (Biolegend, M1/70), anti-CD8-PE (Biolegend, 53-6.7), anti-CD4-BV785 (Biolegend, RM4-5), anti-CD19-BV785 (Biolegend, 6D5), anti-Ly6G-PE (Biolegend, 1A8), anti-NK1.1-PeCy7 (Biolegend, PK136), anti-MHCII-FITC (Biolegend, M5/114.15.2), anti-CD16/32-PerCP-Cy5.5 (BD Bioscience, 2.4G2), anti-SiglecF-PE-Efluor610 (eBioscience, 1RNM44N), anti-Ly6C-AlexaFluor700 9Biolegend, HK1.4), and anti-CD86-APC-Cy7 (Biolegend, GL-1). Following two washes with PBS, the samples were fixed and permeabilized using the Foxp3 staining kit (eBioscience), according to the manufacturer’s protocol. After permeabilization, the samples were incubated with anti-CD68-BV421 (BD Bioscience, FA/11) for 30 minutes at 4 °C. Samples were acquired on an Aurora full spectrum cytometer (Cytek) in the Cincinnati Children’s Hospital medical center Flow cytometry core. Unstained and single-stained samples were acquired and used to calculate the unmixing parameter using Spectroflow (Cytek). Unmixed samples files were exported as FCS files and analyzed using FlowJo software (BD Bioscience).

Spleen: Mice spleens were extracted and minced into 1mm pieces using a razor blade. The minced hearts were washed three times with phosphate-buffered saline (PBS), to remove intravascular cells. The Splenocytes were then digested for 45 minutes at 37 °C under agitation, in Hank’s Balanced Salt Solution (HBSS) solution containing 1mg/ml Collagenase IV (Gibco) and 10ug/ml DNase I (Sigma-Aldrich). Cell suspensions were filtered through a 70µm strainer, and debris were removed using Debris removal solution (Miltenyi), following the supplier’s protocol. For immunophenotyping by flow cytometry, two million splenocytes per sample were used. The single cell suspensions were first labeled for 30 minutes at 4 °C with LIVE/DEAD Fixable Blue (Invitrogen), to separate live cells from dead cells during analysis. After two washes, the samples were incubated for 30 minutes at 4 °C with the following antibodies: anti MO CD4 Brilliant violet 500 (eBioscience RM 4-5), anti Mo CD8a-Super Bright 702 (eBioscience 53-6.7), anti MO CD163-Super Bright 600 (eBioscience TNKUPJ), anti FoxP3 Alexa Fluor 488 (eBioscience 150D/E4), anti Helios-PE (eBioscience 22F6), anti Mo CD16/32-PerCP cy5.5 (eBioscience 93). Following two washes with PBS, the samples were fixed and permeabilized using the Foxp3 staining kit (eBioscience), according to the manufacturer’s protocol. After permeabilization, the samples were incubated with anti MO CD68-Brilliant violet 786 (eBioscience FA-11) for 30 minutes at 4 °C. Samples were acquired on AttuneNXT (Thermo Fisher Scientific). Unstained and single-stained samples were acquired and analyzed using FCS express (De Novo software).

### Immunoblotting for Intestinal Tight Junction Proteins

About 5 mm of tissue from the large intestine prior to cecum was dissected and flash frozen. Total protein was extracted by homogenization in RIPA buffer supplemented with HaltTM protease inhibitor cocktail. Protein levels were normalized, and samples were loaded into a 12% tris-glycine gel and SDS-PAGE was performed in a tris/glycine/SDS buffer system (Biorad, Hercules, CA). The gel was transferred to a 0.45 µm PVDF membrane, blocked in 5% milk-TBS-Tween and incubated with primary antibodies (Proteintech, Rosemont, IL) at manufacturer recommended dilutions for 2 hours. Membrane was washed three times with TBS-Tween, followed by incubation with relevant HRP-conjugated secondary antibody (Proteintech). After three subsequent washes with TBS-Tween, the membrane was developed using SupersignalTM West Pico PLUS Chemiluminescent Substrate (Thermo Scientific, Waltham, MA). The blot was detected using the C-DiGit Blot Scanner (LI-COR Biosciences, Lincoln, NE).

### ELISA

ELISA for serum lipopolysaccharide binding protein was performed with mouse serum using Mouse LBP ELISA Kit PicoKine^TM^ (BosterBio, Pleasanton, CA) as per manufacturer’s instructions. Serum was obtained by allowing mouse blood to clot at room temperature for 30 min followed by refrigerated centrifugation at 2000xg for 15 min.

ELISA for CRP was performed using Mouse C-Reactive Protein/CRP ELISA Kit from ProteinTech, Rosemont, IL as per manufacturer’s instructions using mouse serum.

### PRIMERS

Primers used in the study can be found below (5’-3’)

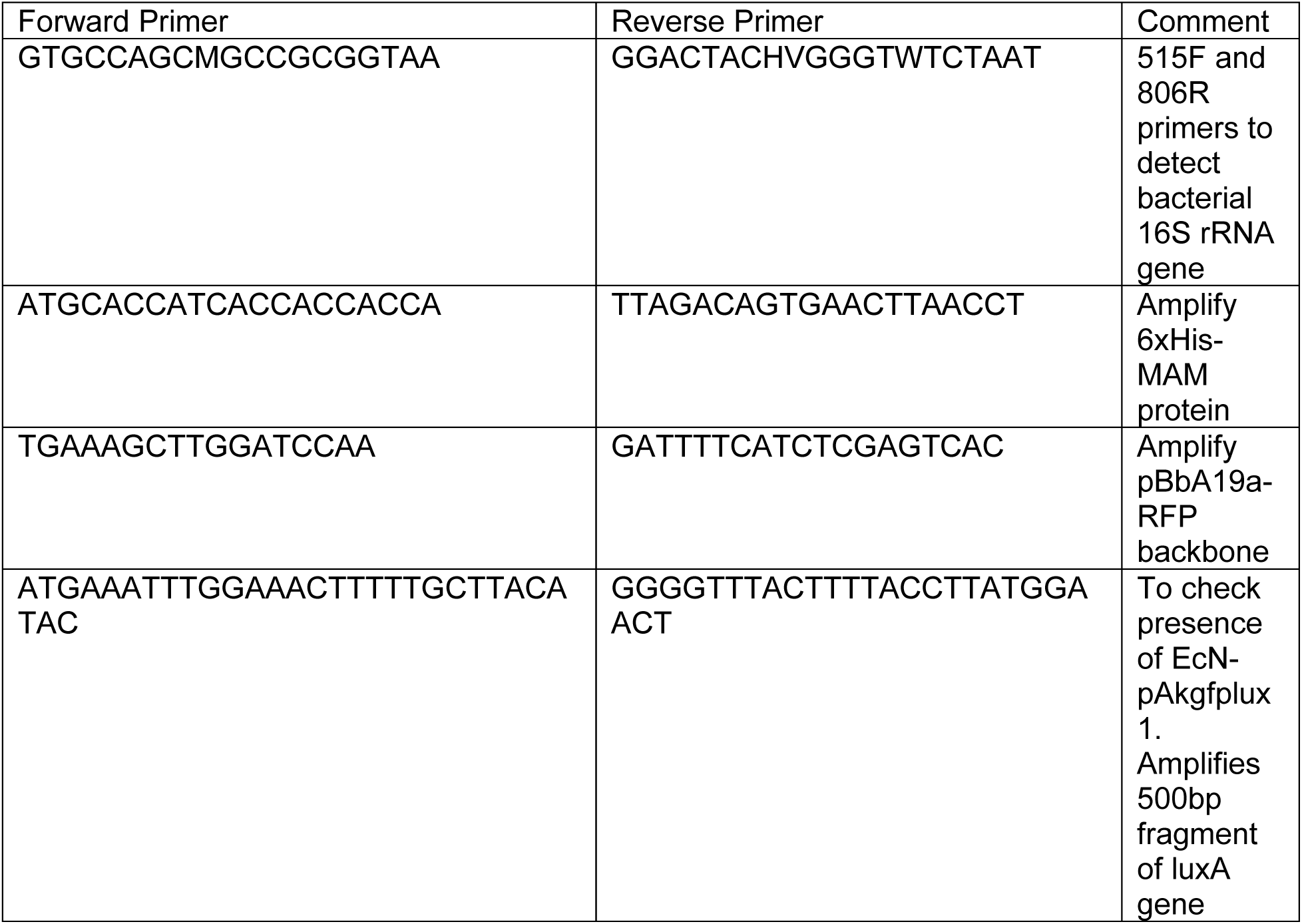

### BACTERIA

Bacteria used in the study are as follows

BACTERIA Bacteria used in the study are as follows
| Bacteria | Strain Abbreviation | Plasmid | Comment |
| --- | --- | --- | --- |
| *Escherichia coli*<br>Nissle 1917 | EcN-Lux | pAKgfplux1 | *Escherichia coli*<br>Nissle 1917<br>expressing luciferase<br>and GFP |
| *Escherichia coli*<br>Nissle 1917 | EcN-MAM | pBbA19a-MAM | *Escherichia coli*<br>Nissle 1917<br>expressing MAM<br>protein derived from<br>*Fecalibacterium*<br>*prausnitzii* |

### STATISTICAL ANALYSES

Data has been represented either as mean ± SD or mean ± SEM and has been indicated accordingly in the figure legends. Sample size for each experiment has also been indicated in the figure legends. Ordinary one-way ANOVA (Tukey’s multiple comparison test) or unpaired t-test were used as indicated, to compare means between groups. Values of p<0.05 were demed to be statistically significant. Statistical analyses was performed either using GraphPad Prism 8 software or R.

## ACKNOWLEDGEMENTS

The authors acknowledge the help of Xiangning Wang and Dr. Lisa Lemen from the University of Cincinnati College of Medicine Imaging Core. We would also like to thank Cincinnati Children’s Hospital Medical Center Pathology Department. Dr. Greg Caporaso and his research group for support related to microbiome analysis using QIIME2. This work was supported by grants from NIH: R01HL168588 and R01CA279962, and American Heart Association: AHA 24TPA1303687 (to N.K.), NIH: R01AR078001, R01AR079435, R01AR079477, R01HL105826, R01HL130356 and AHA: 19TPA34830084, 945748 and Institutional Award for Undergraduate Student Training: 24IAUST1194774 (to S.S.), and Research Fellowship Award from the University of Cincinnati Graduate School (to T.M., S.C.T., N.S.).

## DATA AVAILABILITY

The data that support the findings of this study are available from the corresponding author upon reasonable request.

## MATERIALS AVAILABILITY

MAM plasmid generated in this study will be made available on request, but we may require a a completed materials transfer agreement if there is potential for commercial application.

## AUTHOR CONTRIBUTIONS

T.M. and N.K. conceived the project and designed the experiments. T.M. designed and cloned the constructs. T.M, M.L.N., D.G., A.H., M.S., N.S.K., S.C.T., S.E.K., and L.K. conducted the animal experiments. T.M. and N.S. conducted PET/CT imaging. T.M., M.S., and N.S.K. conducted flow cytometry analysis. T.M. and T.R. conducted microbiome analyses. S.S. and J.L.N. provided input about experimental design. T.M. and N.K. analyzed the data and wrote the manuscript with feedback and review from all the authors.

## SUPPLEMENTARY DATA

**Supplementary Fig. 1.**
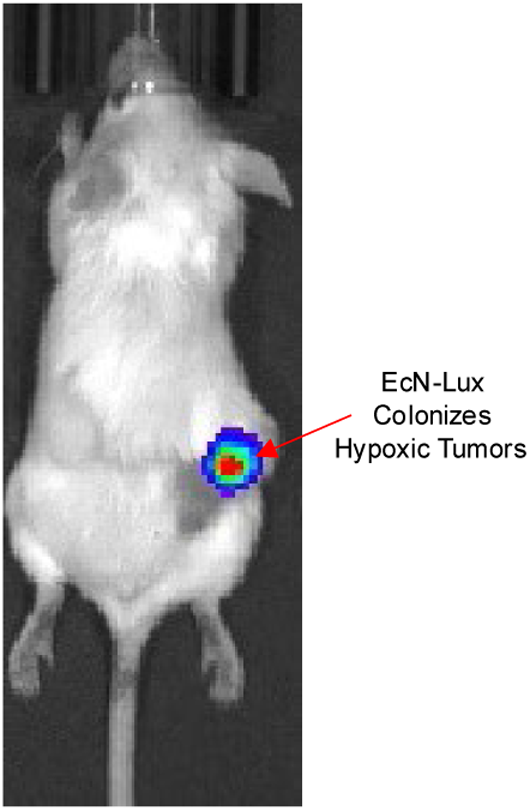
(A) Bioluminescence imaging shows colonization of intravenously administered EcN-Lux in subcutaneous tumor

**Supplementary Fig. 2.**
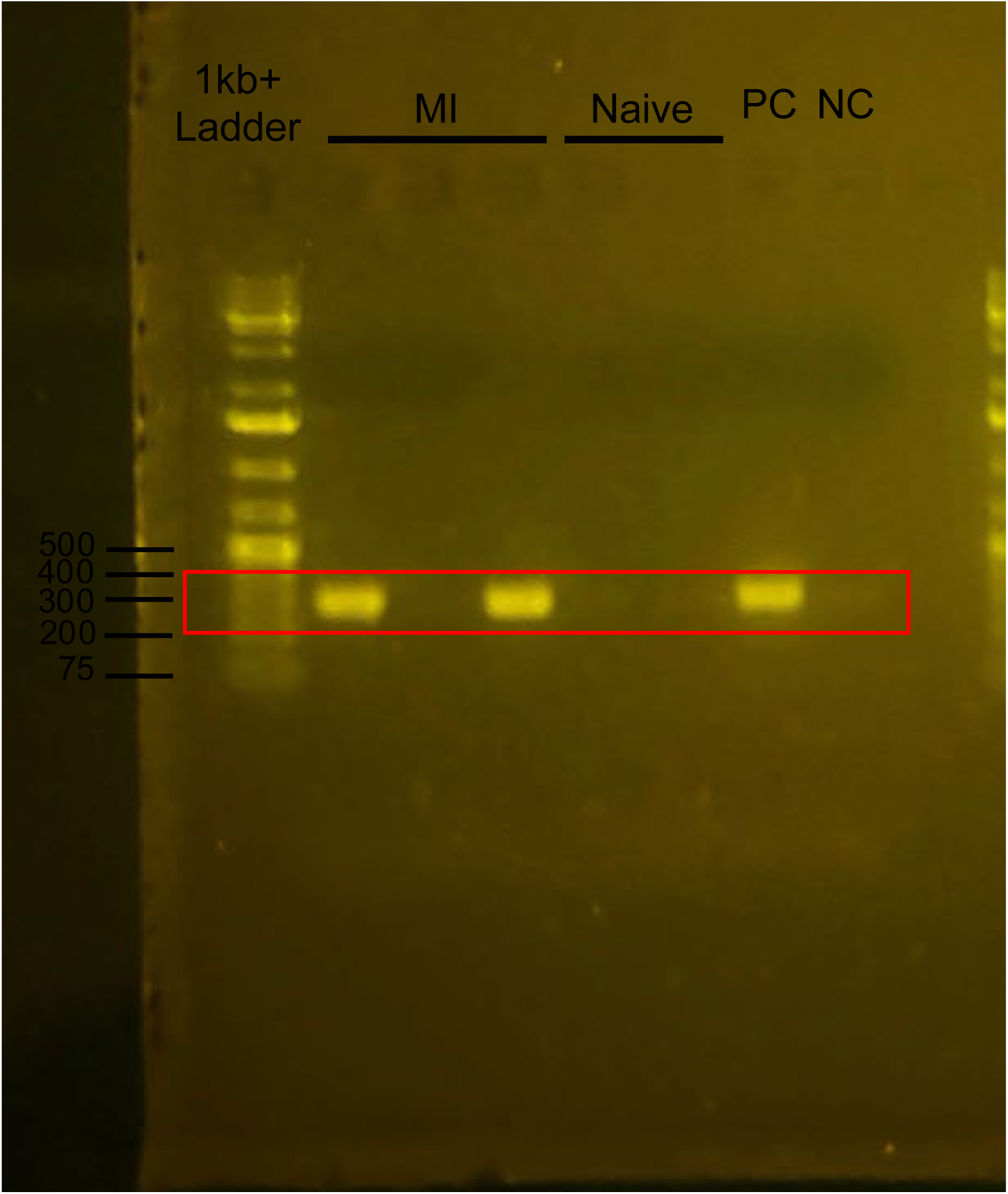
(A) PCR for 16S rRNA shows presence of bacterial DNA in hearts of pigs that underwent IRI surgery (n=3), but not naïve pigs (n=2). Ladder markers are in base pairs (bp).

**Supplementary Fig. 3.**
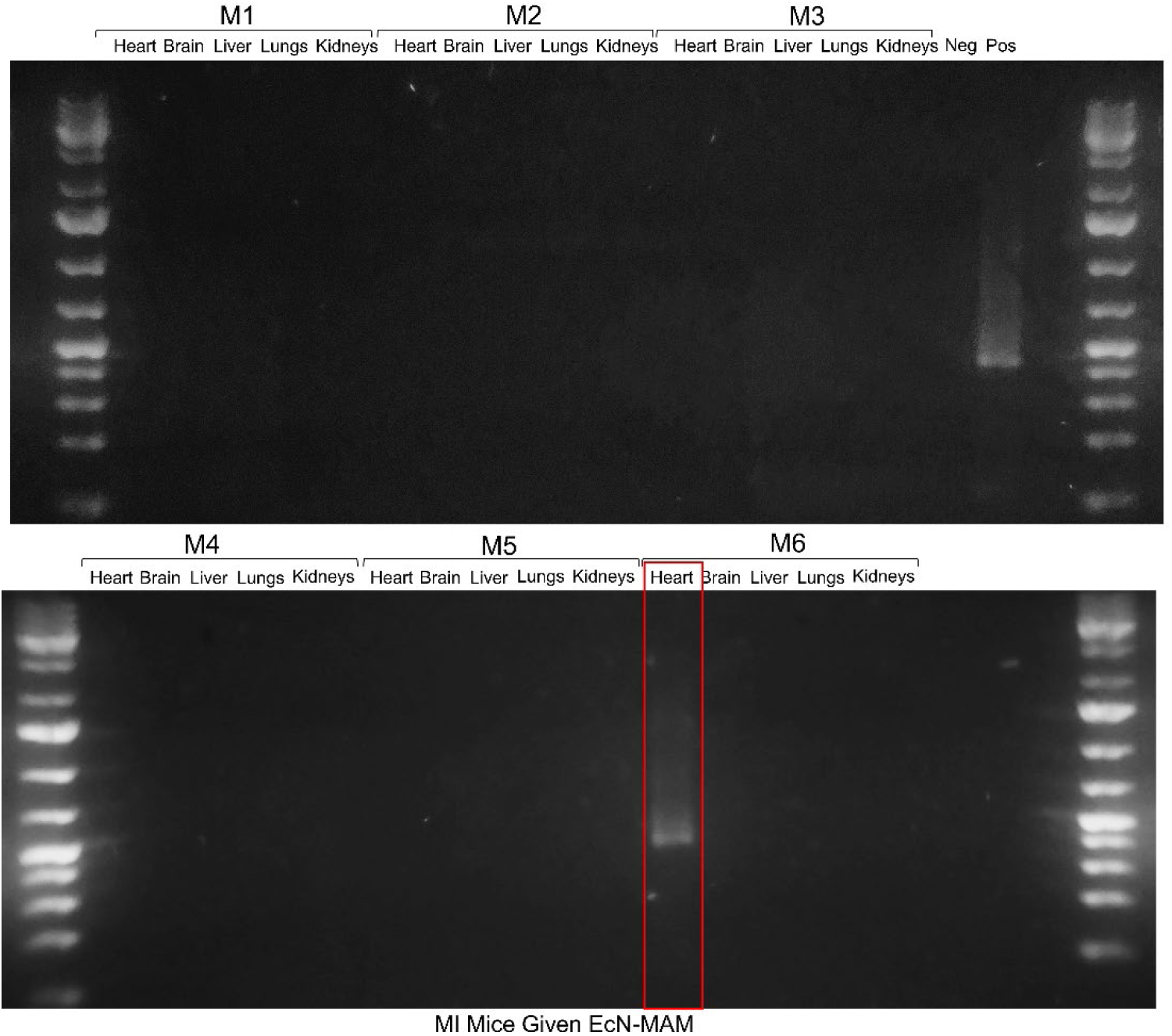
PCR of major organs of EcN-MAM-treated MI mice revealed that EcN-MAM does not distribute or colonize any major organs.

**Supplementary Fig. 4.**
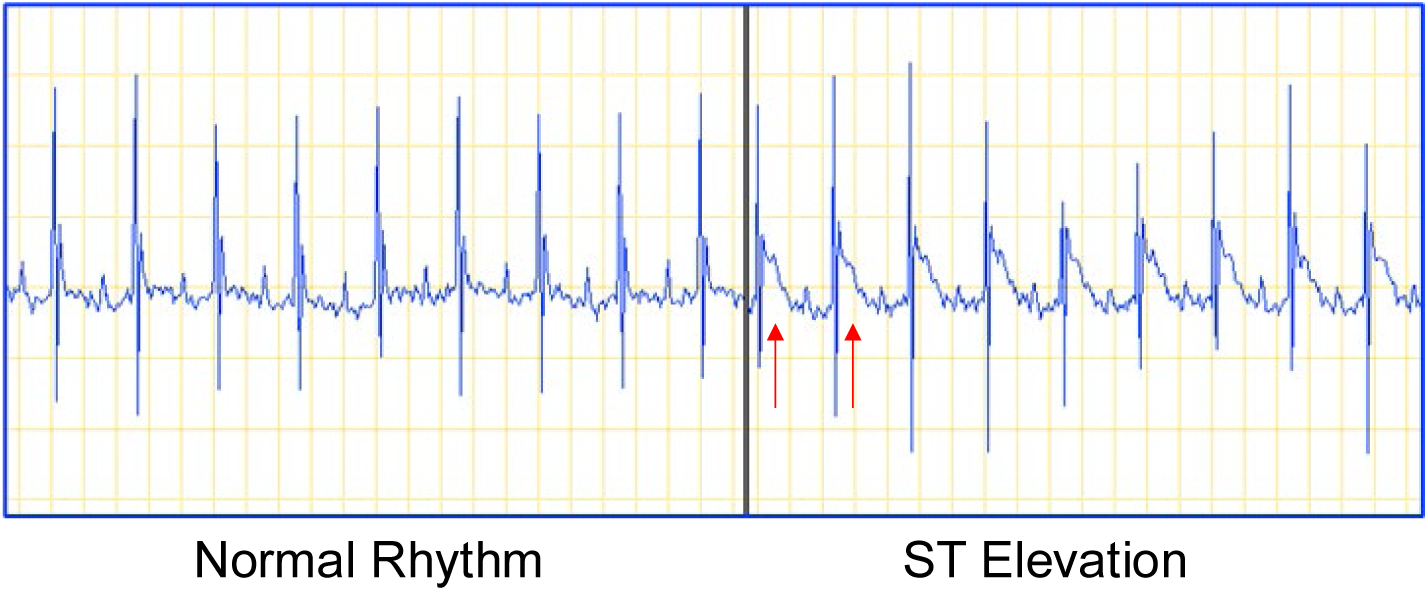
Occlusion of left anterior descending (LAD) artery was confirmed by observing elevation of ST segment in the electrocardiogram.

**Supplementary Fig. 5.**
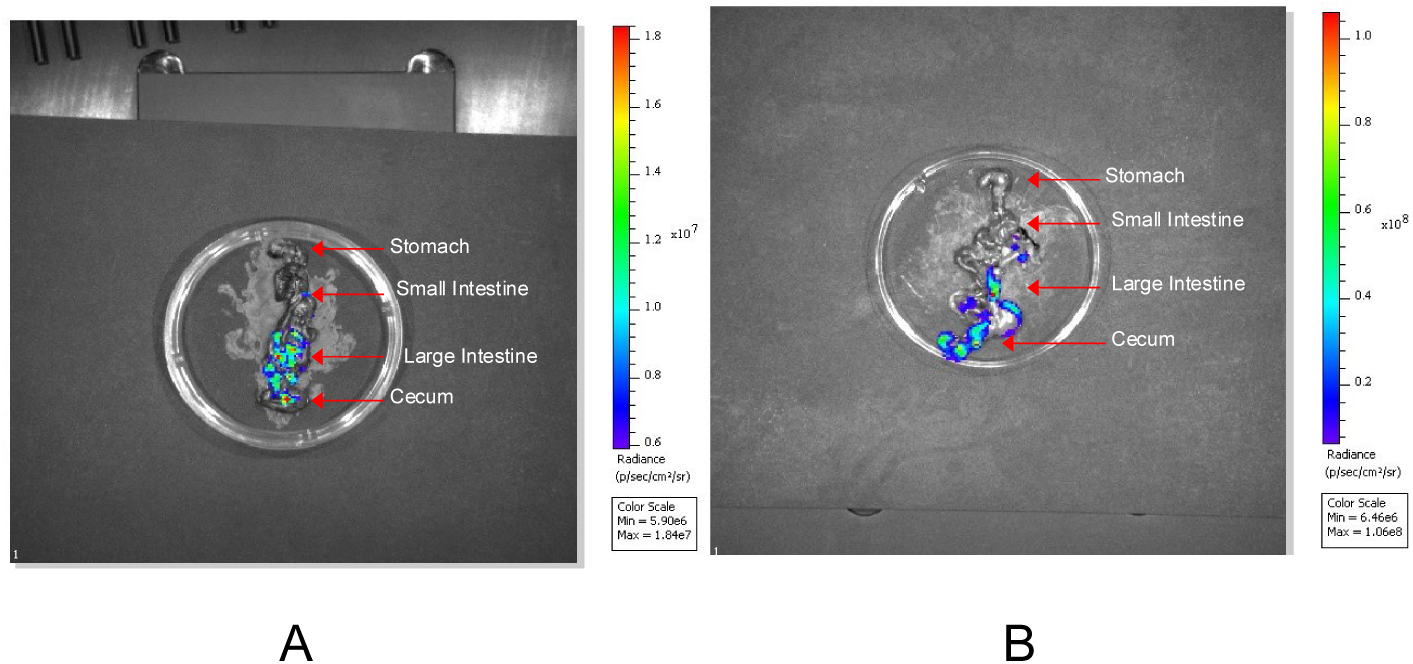
(A-B) Ex vivo IVIS imaging of GI tract indicates EcN Lux primarily colonizes distal region of the intestines, particularly colon and cecum at 48h (A) and 1-week (B) timepoints.

